# Complex-Traits Genetics Virtual Lab: A community-driven web platform for post-GWAS analyses

**DOI:** 10.1101/518027

**Authors:** Gabriel Cuéllar-Partida, Mischa Lundberg, Pik Fang Kho, Shannon D’Urso, Luis F. Gutiérrez-Mondragón, Trung Thanh Ngo, Liang-Dar Hwang

## Abstract

**Motivation:** For over a decade, genome-wide association studies (GWAS) have been an important method for mapping genetic variation underlying complex traits. With the ever-increasing volume of data generated however, new tools are needed to integrate the vast array of GWAS results and perform further analyses in order to maximize their utility for biological discovery. Here we present the Complex-Traits Genetics Virtual Lab (CTG-VL) — https://genoma.io.

**Results:** CTG-VL integrates several key components: **(i)** publicly available GWAS summary statistics; **(ii)** a suite of analysis tools; **(iii)** visualization functions; and **(iv)** data sets for genomic annotations. The platform also makes available results from gene-, pathway- and tissue-based analyses of >1,500 complex traits for assessing pleiotropy at the genetic variant through to these higher levels. Here we demonstrate the platform by re-analysing GWAS summary statistics of back pain (N=509,070). Using analysis tools in CTG-VL we identified 59 genes, of which 20 are across 10 loci outside the original GWAS signals — including *NTRK1* — which is important for the development of pain-mediating sensory neurons. Further, we found enrichment for a number of central nervous system regions in back pain, and evidence for a potential causal relationship with height (OR = 1.06 per cm; 95%CI = 1.04 – 1.08). Using CTG-VL’s database, we show biological pathways associated with back pain are also associated with other traits such as self-reported temperament (‘highly strung’), walking and mood swings.

**Conclusions:** CTG-VL is a freely available online web application to further harness GWAS data for research reproducibility, collaboration and translation.

## Background

Genome-wide association studies (GWAS) have revolutionized genetics research of complex traits and diseases over the past decade (Cloney, 2016; Visscher, et al., 2017). As these studies become more common thanks to initiatives such as UK Biobank (Sudlow, et al., 2015) and many international genetics research consortia, there is a growing need for large-scale data sharing and readily accessible collaborative genomic analyses tools for users with varied technical backgrounds.

Toward this end, we present the first public release of the Complex-Traits Genetics Virtual Lab (CTG-VL) — a freely available web platform to annotate, analyse and share GWAS/post-GWAS results. CTG-VL complements already widely used web applications such as LD-Hub (Zheng, et al., 2017), FUMA-GWAS (Watanabe, et al., 2017) and MR-Base (Hemani, et al., 2018) by incorporating functions available in these platforms along with many others. In this paper, we compare CTG-VL with these other web applications and demonstrate the platform’s utility by applying a range of its functions to GWAS summary statistics of back pain (Freidin, et al., 2019).

## Results

### Visualization and annotations

In the current release of CTG-VL, visual inspection of GWAS results is through LocusTrack plots (Cuellar-Partida, et al., 2015) and Manhattan plots. These visualizations are fully interactive where users can annotate and query results with a single click. For annotation, we incorporated epigenetic data from ENCODE (Consortium, 2012) and Roadmap epigenomics projects (Bernstein, et al., 2010), eQTL data from the GTEx project (Consortium, 2013), GWAS catalogue (MacArthur, et al., 2017) and polyphen (Adzhubei, et al., 2013). Other utilities such as obtaining the closest genes for a given genetic variant, obtaining gene information, SNPs in linkage disequilibrium (LD) and population allele frequencies, are also actionable through a single click. Furthermore, CTG-VL provides other graphing/plotting functions such as heatmaps, karyograms and network visualizations. For the latter, we incorporated functions to analyse the topology of networks including density, degree distribution, mean degree, number of edges of the giant component, as well as functions to estimate the shortest paths between nodes.

### Analysis tools

The goal of CTG-VL is to facilitate downstream analyses of GWAS summary statistics. As such, we are actively integrating analysis tools and data that will enable the research community to speed up their analyses. The platform’s current release (0.39-alpha) includes a selection of the newest and most commonly used analysis tools for downstream analyses of GWAS results. Briefly, we have integrated LD-score regression (Bulik-Sullivan, et al., 2015; Bulik-Sullivan, et al., 2015) to estimate the heritability of traits and genetic correlation between traits using GWAS summary statistics. Users can also check genetic correlations between any of the >1,500 traits with publicly available GWAS summary statistics integrated into the platform.

CTG-VL implements Data-driven Expression-Prioritized Integration for Complex Traits (DEPICT) (Pers, et al., 2015) — a tool that prioritizes the most likely causal genes at associated loci, and identifies enriched pathways and tissues/cell types which may underlie the associations. Another popular tool that has been incorporated is MetaXcan (Barbeira, et al., 2018), which obtains gene-trait association by testing if the predicted expression levels of particular genes (based on prediction models derived from eQTL data of selected tissues) underlie the GWAS associations. Similarly, we integrated fastBAT (Bakshi, et al., 2016) — a fast set-based association analysis that uses GWAS summary statistics and a linkage-disequilibrium reference to summarize genetic associations with a trait of interest at the gene level. Finally, we have included Summary-data-based Mendelian Randomization (SMR) (Zhu, et al., 2016) to test the causal effect of gene expression levels on a trait of interest, and Generalized Summary-data-based Mendelian Randomization (GSMR) (Zhu, et al., 2018) to test the causal relationship between two traits.

### Data

CTG-VL is an aggregator of GWAS summary statistics and post-GWAS analyses results. It therefore complements the recent release of GWAS-ATLAS (Kyoko Watanabe, 2018) — a titanic effort that integrates results from post-GWAS analyses of >3,500 complex traits in a browsable platform (e.g., gene-based and gene-set enrichment analyses results from MAGMA (de Leeuw, et al., 2015) — along with genetic correlations and heritability derived from LD-score and SumHer (Speed and Balding, 2018)). CTG-VL has made available post-GWAS results of >1,500 traits (with DEPICT and MetaXcan) and we are actively updating this resource. CTG-VL also enables users to readily upload their own GWAS summary statistics or to link the >1,500 GWAS summary statistics integrated in the platform with their user profile and perform further analyses.

### Web applications ecosystem

Comparing CTG-VL with other currently available web platforms (**Table 1**) can be challenging as they each facilitate different specific tasks and are optimized for particular pipelines/analysis methods. While all these tools aggregate GWAS summary statistics, their functions are largely segregated. CTG-VL complements other web applications already available, by integrating their key analysis methods into a single web platform. **Table 1** outlines key differences and similarities between CTG-VL and three widely used web tools: FUMA-GWAS, LD-Hub and MR-Base.

**Table 1.** A comparison of features in CTG-VL, FUMA-GWAS, LD-HUB and MR-Base.

|  | CTG-VL | FUMA-GWAS | LD-Hub | MR-Base |
| --- | --- | --- | --- | --- |
| **Aggregator of GWAS summary statistics** | Yes | Yes | Yes | Yes |
| **Visualization** | All visualizations are interactive:<br>Manhattan plot<br>Regional plot<br>Heatmaps<br>Network plots<br>Karyograms | Manhattan plot<br>Regional plot (interactive)<br>Heatmap (only for gene expression) | No | No |
| **Data for annotation** | ENCODE<br>Roadmap Epigenomics<br>GTEx<br>Polyphen | ENCODE<br>Roadmap Epigenomics<br>GTEx<br>Polyphen<br>Multiple others<br>(http://fuma.ctglab.nl/links) | No | No |
| **Heritability** | LD-score regression | No | LD-score regression | No |
| **Genetic correlations** | LD-score regression<br>User chooses trait of interest from available GWAS in the platform (>1500 complex traits) or their own GWAS dataset.<br>Automated to run against >1500 complex traits if the user chooses to. | No | LD-score regression | No |
| **Gene-based analysis** | MetaXcan<br>fastBAT<br>SMR<br>DEPICT gene-prioritization | MAGMA | No | No |
| **Tissue specificity analysis** | DEPICT | MAGMA | No | No |
| **Gene-enrichment analysis** | DEPICT | MAGMA | No | No |
| **Mendelian Randomization** | GSMR<br>SMR | No | No | Over 10 different statistical methods to perform two-sample Mendelian Randomization (excluding SMR & GSMR). |
| **Meta-analysis** | MTAG | No | No | No |

### Case study

In this section, we illustrate a series of analyses that can be performed in CTG-VL. Summary statistics from a recent GWAS meta-analysis study of back pain by Freidin et al. (2019) were used as an example. This back pain dataset along with >1500 other traits are available in CTG-VL. **Figure 1** outlines the key analysis tasks performed in the platform. Screenshots capturing each step are shown in **Supplementary Figures 1–9**. Using the tools in CTG-VL, we generated LocusTrack and Manhattan plots (**Figures 2 & 3**). Like Freidin et al. (2019), we performed pathway and tissue enrichment analyses with DEPICT and computed genetic correlations using LD-score regression. We also assessed causal relationships of gene expression and risk factors on back pain using SMR and GSMR, respectively. Next, we performed gene-based analyses with fastBAT and used CTG-VL’s large database to identify other complex traits that have shared biological pathways with back pain.

**Figure 1.**
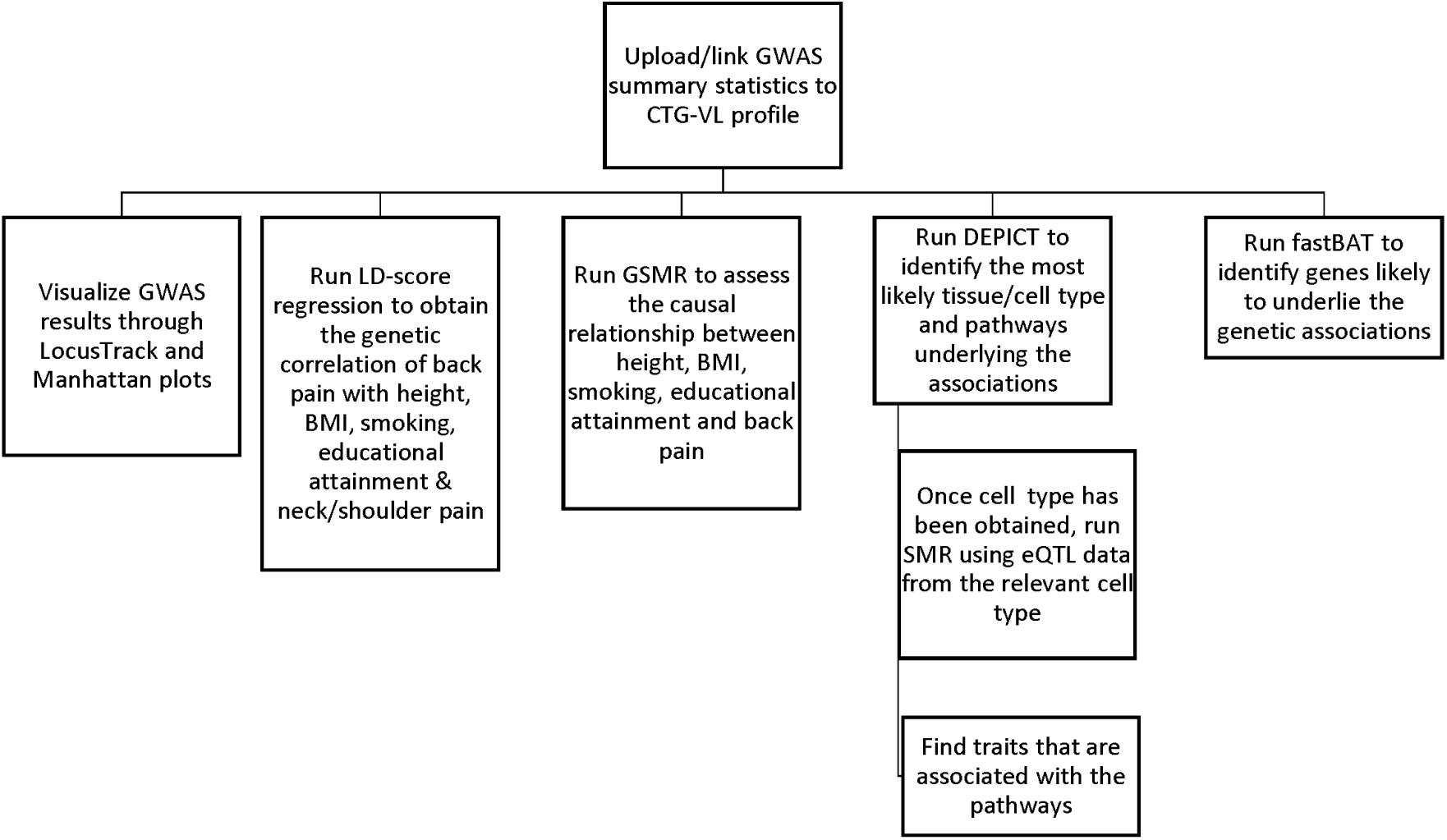
Key tasks in the re-analysis of back pain GWAS summary statistics using CTG-VL

**Figure 2.**
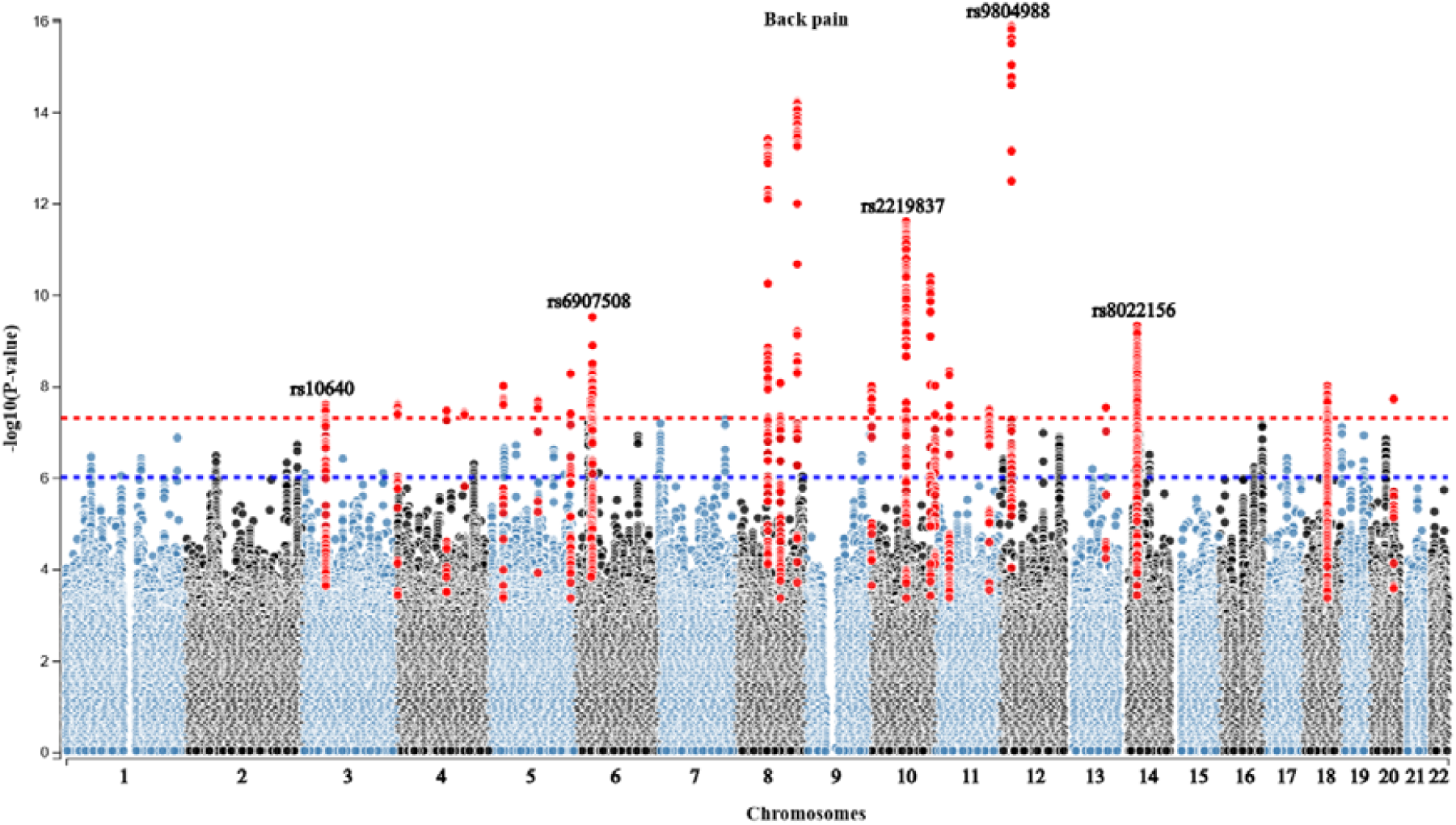
Manhattan plot of back pain post-GWAS analysis. CTG-VL enables the user to click on each of the SNPs and obtain the summary statistics as well as annotate and query the SNP. The user can also highlight SNPs in LD with the lead SNP, as shown in red.

**Figure 3.**
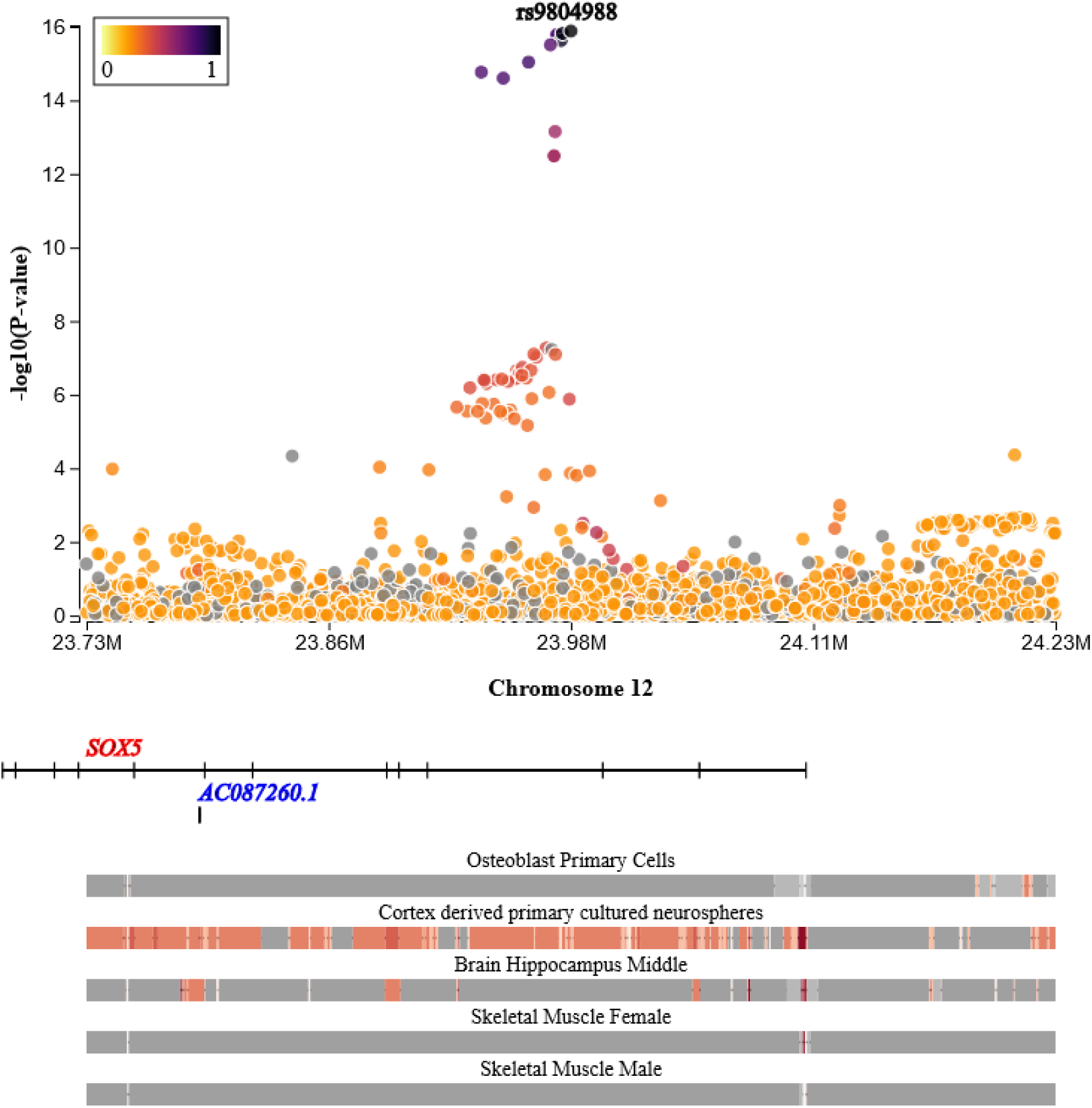
LocusTrack plot of the top GWAS hit for back pain. CTG-VL enables the user to generate regional plots of any SNP from the GWAS results. The plot is fully interactive and allows plotting annotation tracks, which are shown here and correspond to chromatin state annotation for diverse tissues. Red intensity in the annotation track indicates open chromatin/active transcription.

The first step in carrying out these analyses was to link the relevant data to our user profile, e.g., *Data → Public data* (**Supplementary Figure 1**). Here, the user selects the relevant GWAS summary statistics necessary for the analyses and links them to their profile by clicking *“Add to my data”*. In our particular example, we were interested in estimating the genetic correlation of back pain (Freidin, et al., 2019) with height, BMI, ‘ever smoked’, neck/shoulder pain (Lab, 2018) and educational attainment (Lee, et al., 2018), which were added to “my data”. The back pain GWAS results were then visualised by generating a Manhattan plot (*Visualization → Manhattan plot*) and a LocusTrack plot (*Visualization → LocusTrack*) of the loci with the strongest association (**Supplementary Figures 2 & 3**). CTG-VL implements interactive Manhattan and LocusTrack plots, which enable the user to annotate the SNP of interest by clicking on it. In addition, users can highlight all the SNPs that are in LD with the lead SNP of each locus.

We estimated the genetic correlation (*Analyses → LD-score*) of back pain with BMI, height, ‘ever smoked’, educational attainment and neck/shoulder pain (**Supplementary Figure 4**). In line with the findings by Freidin *et al*. (2019), we observed: **(i)** statistically significant positive genetic correlations between back pain and BMI (r_g_ = 0.34 ± 0.02; P = 1.02 ×10^−63^), smoking (r_g_ = 0.23 ± 0.02; P = 2.05 ×10^−16^) and neck/shoulder pain (r_g_ = 0.72 ± 0.11; P = 8.18 ×10^−11^); **(ii)** a statistically significant negative genetic correlation between back pain and educational attainment (r_g_ = −0.45 ± 0.02; P = 2.48 ×10^−105^); and **(iii)** a null genetic correlation between back pain and height (r_g_ = 0.01 ± 0.02; P = 0.68), showing that the platform generates reproducible results.

Using GSMR(Zhu, et al., 2018) in CTG-VL (*Analyses → GSMR*), we examined whether height and BMI had a causal effect on back pain (**Supplementary Figure 5**). GSMR is a two-sample Mendelian Randomization approach, which requires GWAS summary statistics from the outcome (e.g., back pain) and exposures (e.g., height & BMI) to be derived from independent samples (Zhu, et al., 2018). Therefore, we used GWAS summary statistics for height and BMI in previous studies from the Genetic Investigation of ANthropometric Traits (GIANT) consortium that did not include UK Biobank data (Locke, et al., 2015; Wood, et al., 2014). We observed evidence for a causal effect of height on back pain (OR = 1.06 per cm increase in height; 95% CI = 1.04 – 1.08), but not for BMI on back pain (OR = 1.04 per kg/m^2^; 95% CI = 0.99 – 1.09). However, the Heterogeneity In Dependent Instruments (HEIDI) test for these analyses were marginally significant (P_HEIDI_Height_ = 0.057 and P_HEIDI_BMI_ = 0.0504), suggesting the presence of some instrumental variables (SNPs) with pleiotropic effects (i.e., the SNPs may be affecting the outcome not through the exposure of interest).

Next, we ran DEPICT (*Analyses → DEPICT*) to obtain the most likely tissues/cell types and pathways underlying the GWAS associations (**Supplementary Figure 6**). **Table 2** presents the tissues and cell types we found to be associated with back pain. DEPICT takes independent SNPs below a specific significance threshold as input SNPs. We used the same thresholds as Freidin *et al*. (P<1.0 ×10^−5^ and P<5.0 ×10^−8^). In line with their findings, we did not observe any significant (FDR < 5%) tissue/cell type association using independent SNPs at P<5.0 x10^−8^. In contrast however, we identified associated tissues (FDR < 5%) when using independent SNPs at P<1.0 x10^−5^. This difference in results could be attributable to the way independent SNPs were selected. CTG-VL uses clumping — i.e., iteratively removing SNPs correlated to the SNPs with highest p-value within a 500Kb window — as oulined in the DEPICT documentation. The study by Freidin *et al*. (2019) performed conditional analyses to select the independent SNPs. Other factors that could lead to the difference in results is the reference genotype used to identify the independent loci. CTG-VL uses the European population genotype sample from 1000 Genomes, whereas the study by Freidin *et al*. used 10,000 individuals drawn from UK Biobank. Furthermore, CTG-VL implementation ensures all the input SNPs are available in DEPICT’s models, whereas it is not clear if that was checked in their study. Our pathway analyses using DEPICT did not reveal any significant association (FDR < 5%) using either of the two significance thresholds. These results are shown in **Supplementary Table 1**. Comparisons between them and pathway analysis results from the study by Freidin *et al.* are presented in **Supplementary Tables 2 & 3.**

**Table 2.** Tissues and cell types enriched for back pain (FDR < 5%) derived from DEPICT using independent SNPs at P<1.0 ×10^−5^

| Tissue | Group | P-value |
| --- | --- | --- |
| Retina | Sense Organs | 0.000858 |
| Brain | Nervous System | 0.000931 |
| Central Nervous System | Nervous System | 0.000932 |
| Parietal Lobe | Nervous System | 0.00125 |
| Hippocampus | Nervous System | 0.00128 |
| Cerebral Cortex | Nervous System | 0.00174 |
| Prosencephalon | Nervous System | 0.00174 |
| Limbic System | Nervous System | 0.00186 |
| Neural Stem Cells | Cells | 0.00192 |
| Telencephalon | Nervous System | 0.00206 |
| Cerebrum | Nervous System | 0.00213 |
| **Parahippocampal Gyrus** | Nervous System | 0.00288 |
| **Entorhinal Cortex** | Nervous System | 0.00288 |
| **Visual Cortex** | Nervous System | 0.00308 |
| **Occipital Lobe** | Nervous System | 0.00310 |
| **Temporal Lobe** | Nervous System | 0.00352 |

In CTG-VL, we used the “check overlap” function and the large database of post-GWAS analyses on >1,500 complex traits to examine whether other traits and diseases are also associated with back pain top-associated pathways (**Supplementary Figure 7**). We found 9 out of 17 pathways that had a suggestive association with back pain (P < 0.0005) were also associated with self-reported tension (‘highly strung’); 8 pathways were associated with walking (i.e., type of transport used); and 6 pathways were associated with mood swings (**Supplementary Tables 4 & 5**).

Next we carried out SMR analysis (*Analysis → SMR*) using data derived from a meta-analysis of eQTL data on all brain tissues from GTEx (Qi, et al., 2018). This analysis prioritizes genes that are more likely to be associated with back pain (**Supplementary Figure 8**). In total, we examined 7,221 genes and hence the Bonferroni corrected p-value threshold for significance was P < 7e-6. Significant SMR results are shown in **Table 3**.

**Table 3.** Significant SMR results based on eQTL data from all brain tissues (GTEx).

| Gene | CHR | Position | eQTL | Effect | P-value | P-value<sub>HEIDI</sub>* | No. SNPs |
| --- | --- | --- | --- | --- | --- | --- | --- |
| **C6orf106** | 6 | 34609850 | rs3800461 | 0.03 | $2.18 \times 10^{-9}$ | 0.001 | 20 |
| **CHST3** | 10 | 73748722 | rs2091331 | -0.08 | $3.42 \times 10^{-8}$ | 0.35 | 20 |
| **GPX1** | 3 | 49395321 | rs9859556 | 0.02 | $4.34 \times 10^{-8}$ | 0.04 | 20 |
| **RNF212** | 4 | 1078694 | rs750305 | 0.05 | $4.46 \times 10^{-7}$ | 0.65 | 20 |
| **SNRPC** | 6 | 34733377 | rs35839188 | -0.02 | $5.81 \times 10^{-6}$ | 0.002 | 18 |

\*HEIDI (HEterogeneity In Dependent Instruments) test P-value. A P-value < 0.05 indicates that the association may be confounded by SNPs in LD with the eQTL.

**Table 4.** Genes significantly associated with back pain (fastBAT gene-based test) outside the original GWAS leading signals.

| Gene | Chromosome | Position | P-value | Top-SNP | No. SNPs |
| --- | --- | --- | --- | --- | --- |
| **20** | *GSS* | 33516235 | $1.36 \times 10^{-7}$ | rs1475070 | 50 |
| **1** | *SH2D2A* | 156776034 | $1.39 \times 10^{-7}$ | rs11264545 | 76 |
| **1** | *NTRK1* | 156785541 | $1.46 \times 10^{-7}$ | rs11264545 | 128 |
| **20** | *TRPC4AP* | 33590206 | $2.42 \times 10^{-7}$ | rs1475070 | 71 |
| 1 | INSRR | 156810664 | 2.68x10<sup>-7</sup> | rs11264554 | 93 |
| 5 | TRPC7 | 135548998 | 2.89x10<sup>-7</sup> | rs139446184 | 127 |
| 16 | LOC100129697 | 89006446 | 5.62x10<sup>-7</sup> | rs8045567 | 145 |
| 5 | GABRB2 | 160715435 | 5.64x10<sup>-7</sup> | rs7722106 | 210 |
| 20 | MIR499A | 33578178 | 6.63x10<sup>-7</sup> | rs1475070 | 45 |
| 20 | MIR499B | 33578202 | 6.63x10<sup>-7</sup> | rs1475070 | 45 |
| 11 | ZNF408 | 46722316 | 7.23x10<sup>-7</sup> | rs61884303 | 49 |
| 8 | AGO2 | 141541263 | 7.28x10<sup>-7</sup> | rs4498521 | 185 |
| 20 | EDEM2 | 33703159 | 8.63x10<sup>-7</sup> | rs1885115 | 56 |
| 20 | MYH7B | 33543637 | 8.72x10<sup>-7</sup> | rs1475070 | 60 |
| 11 | ARHGAP1 | 46698624 | 0.00000122 | rs61884303 | 54 |
| 6 | LAMA2 | 129204285 | 0.00000124 | rs265361 | 438 |
| 19 | SYMPK | 46318699 | 0.0000014 | rs35318830 | 135 |
| 2 | DIS3L2 | 232826292 | 0.00000146 | rs1453867 | 327 |
| 16 | CBFA2T3 | 88941262 | 0.00000151 | rs8045567 | 241 |
| 20 | ACSS2 | 33462765 | 0.00000196 | rs1475070 | 51 |

Finally, we performed a gene-based test using fastBAT (**Supplementary Figure 9**). The association between 24,097 genes were tested, hence the Bonferroni statistical significance threshold was P<2.0×10^−6^. We identified 59 genes at this significance level, including 20 genes spread across 10 loci that were outside the leading GWAS signals (**Table 4**). All genes significantly associated with back pain along with their description are presented in **Supplementary Table 6.**

## Discussion

We have presented the first release of CTG-VL — a web application integrating the most comprehensive set of post-GWAS analysis tools available into a single platform — which enables users to test and derive novel hypotheses from GWAS summary statistics. A catalogue of GWAS and post-GWAS analysis results for >1,500 complex traits provides users with the ability to explore pleiotropy at the gene, pathway and tissue levels — as we have demonstrated back pain data — and which is currently not available in any other web platforms. This tool and data resource also has considerable utility for the interpretation of genetic correlations. For example, the implemented LD-score regression can be used to assess the extent of genetic correlation. However, such an analysis is unable to implicate the genes and biological pathways underlying the correlation. To address this issue, researchers can use the “*Check overlap*” function in CTG-VL to identify shared genes, tissues and pathways, and thus generate novel hypotheses from the interpretation of genetic correlations.

We used a recent GWAS meta-analysis on back pain (Freidin, et al., 2019) to test the capabilities of CTG-VL. Freidin *et al*. performed a set of analyses that we also implemented in CTG-VL, thus enabling a direct comparison between their results and those obtained from the platform. Our genetic correlation results were consistent with their study, as were our top DEPICT results, although the latter differed according to the significance testing threshold. The difference in DEPICT results could have been due to the approach in selecting independent variables. The study by Freidin *et al*. also used GSMR and SMR, but these methods were employed to assess pleiotropy of the top back pain-associated loci with other traits and gene expression. In contrast, we used GSMR and SMR to assess causality and therefore these sets of results cannot be directly compared. Furthermore, it is important to note that GSMR and SMR analyses should be performed on GWAS summary statistics of independent samples. Deriving instrumental variables (SNPs) for the exposure to perform Mendelian randomization in the same sample as the outcome can lead to a confounded estimate. In their study, certain GSMR analyses used GWAS summary statistics derived from UK Biobank data (for both the exposure and outcome), which may have led to biases in the results.

We also provide additional insights not mentioned in their study, such as results from fastBAT gene-based test. We identified 59 back pain-associated genes, 20 of which were located outside the loci identified in the original GWAS. In particular, interesting associations not previously reported include the *NTRK1* gene, which plays an important role in the development of pain-mediating sensory neurons (Marmigere and Ernfors, 2007), and the *DIS3L2* gene which has been linked to abdominal pain (Kohler, et al., 2019). These results warrant further investigation. We demonstrated the utility of a post-GWAS results database incorporating >1,500 traits by identifying other traits that are also associated with biological pathways in back pain. We envision that as more results are aggregated in the CTG-VL, further novel biological insights and hypotheses will be generated.

This work was inspired by big efforts from multiple research groups making phenome-wide GWAS summary statistics available (e.g., Ben Neale’s lab at the Broad Institute, GeneAtlas (Kyoko Watanabe, 2018) and their released GWAS summary statistics along with SAIGE software (Zhou, et al., 2018)). These rich GWAS results resources await further analysis, and more importantly, interpretation and further empirical investigation. The integration of functions from other GWAS result web platforms into CTG-VL will also be explored (Beck, et al., 2014; Huang, et al., 2018).

The goal of developing CTG-VL into a user-friendly platform is to enable research teams with varying levels of expertise to perform common post-GWAS analyses and aid in the interpretation of this massive flood of data. At the same time, CTG-VL aims to improve research reproducibility and data sharing by enabling researchers to use the same GWAS summary statistics and annotation datasets for performing such post-GWAS analyses. The platform has been made freely available with new analysis tools and data being incorporated as they become available. Its uptake and feedback from genetics researchers, bioinformaticians and health professionals will directly contribute to further refining and enhancing the utility of CTG-VL for the growing open science community.

## Methods

### Implementation

The web interface and server are written in Angular 6 and Node.js, respectively. Scripts to analyze networks were written with python (2.7) using NetworkX (1.8.1) and Igraph (0.7.1) libraries. Tabix (Li, 2011) is used to index and query user GWAS data. Graphics were generated with in-house d3.js v5 scripts. Genomic annotations and user data are stored at a server in the Queensland Research & Innovation Services Cloud. We use Google’s firebase for authentication and meta-data storage. All communications between CTG-VL’s back-end and front-end are encrypted through SSL (Secure Sockets Layer).

### Analysis software

CTG-VL uses PLINK (Chang, et al., 2015) to clump and calculate LD between SNPs. These two functions are mainly to obtain LD data for the regional plots (LocusTrack), identify SNPs in LD with lead SNPs (i.e., those with an association P-value <5e-8), and identify independent loci to be used in DEPICT analyses.

MetaXcan (Barbeira, et al., 2018) is implemented in the platform with the default options, along with gene expression prediction models from PredictDB (Pers, et al., 2015) to carry out gene-based analyses. LD-score regression (Bulik-Sullivan, et al., 2015; Bulik-Sullivan, et al., 2015) is implemented along with LD-scores derived from 1000 Genomes (Bulik-Sullivan, et al., 2015) to estimate heritability and genetic correlations based on GWAS summary statistics. Prior to LD-score analyses, GWAS summary statistics are run through the munge.py utility and merged to HapMap SNPs to ensure that alleles match across different datasets.

DEPICT (Pers, et al., 2015) is implemented to perform gene prioritization, gene enrichment and identify the tissue/cell type most likely to underlie GWAS associations. DEPICT takes independently associated SNPs below a user-specified P-value threshold as input.

SMR (Zhu, et al., 2016) is implemented along with eQTL data for 48 cell types derived from GTEx and downloaded from the SMR website. GSMR and fastBAT (Bakshi, et al., 2016; Zhu, et al., 2018) are functions in GCTA (Yang, et al., 2011) and were implemented with the default parameters.

Each of the individual commands used to run all the analyses included in CTG-VL are displayed in the platform.

### Data

Currently, genotypes from 1000 Genomes phase 3 (Genomes Project, et al., 2015) are used throughout the platform as a reference panel for LD. eQTL data from the GTEx Project (Consortium, 2013) are used for annotation in regional plots, SMR and MetaXcan. Chromatin states (Ernst and Kellis, 2017) derived from ENCODE (Consortium, 2012) and Roadmap Epigenomics Mapping Consortium [9] are used to annotate regional plots. Genes and SNPs position information is based on the GRCh37 human genome assembly. Descriptions of genes are extracted from RefSeq (O’Leary, et al., 2016) and UniProt (UniProt, 2008).

GWAS summary statistics integrated within the CTG-VL platform are updated regularly, and both the references and links can be found in the platform. GWAS summary statistics are integrated as published by the authors, with the exception of summary statistics derived from the Neale lab (Lab, 2018), where we removed SNPs with a MAF < 0.001 or Minor Allele Count < 25 in Cases (for case-control GWAS), as recommended in their documentation.

### Post-GWAS analyses in >1500 traits

We downloaded 3,670 GWAS summary statistics from the Neale Lab (Lab, 2018) and removed variants as described above. LD-score regression was run against each of these GWAS summary statistics. Those traits with a statistically significant heritability at P-value <0.05 (1,747) were selected for further analyses using DEPICT and MetaXcan. In this first public release, we ran MetaXcan using whole-blood gene-expression prediction models as they were derived from a larger sample compared to other tissues. MetaXcan analysis results using gene prediction models for other cell types will be included in the near future. To run DEPICT, we clumped the summary statistics with the recommended parameters (LD-window 500Kb; R^2^ < 0.05) and used SNPs with the recommended P-value threshold of <1e-5. For traits with <10 independent loci, we relaxed the threshold to P-value <1e-4.

### Updates

CTG-VL is updated on a weekly basis and this report may not list/reference all the tools and data in the latest online version. However, this information can be found on the platform’s home page — https://genoma.io. Further post-GWAS analyses are being performed with MetaXcan in other tissues/cell types, SMR and fastBAT, and will be included in future updates of CTG-VL.

## Acknowledgements

GCP is supported by an Australian Research Council Discovery Early Career Research Award [DE180100976]. ML is supported by a Research Training Program Scholarship from The University of Queensland. PFK is supported by a Research Training Program Scholarship from Queensland University of Technology and QIMR Berghofer Medical Research Institute.

## Contributions

GCP conceived and developed the platform, implemented workflows, built databases and performed analyses. ML aggregated GWAS summary statistics and performed analyses. PFK implemented workflows. LFGM implemented scripts for network analysis. SDU performed analyses. LDH collated annotation data and implemented workflows. TTN & GCP conceived and coordinated the application of genetic analysis methods to pain data. All authors contributed to the writing and reviewing of this manuscript.

