## Supplementary Figure for "Complex-Traits Genetics Virtual Lab: A community-driven web platform for post-GWAS analyses"

† Co-last authors

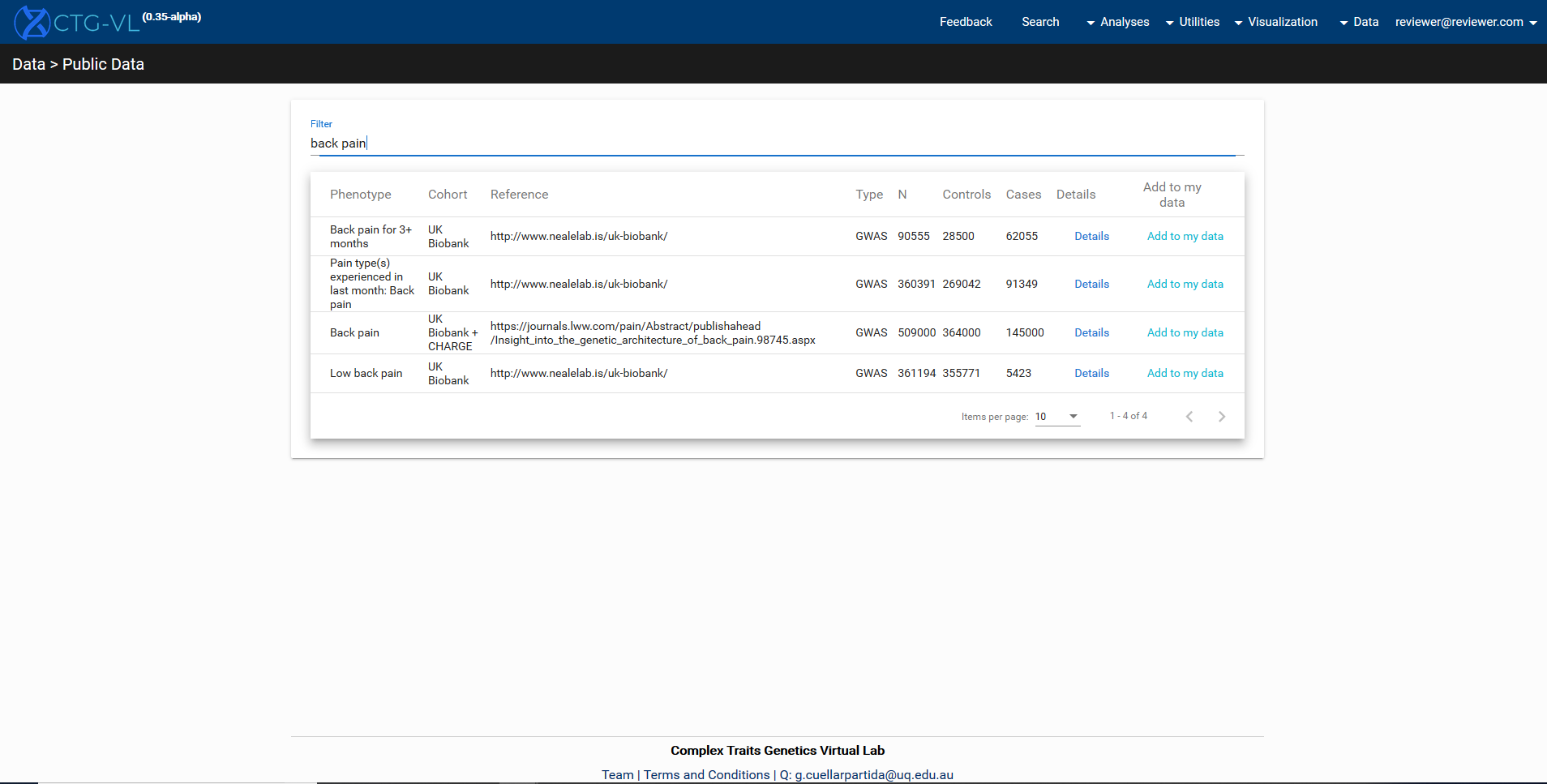

**Supplementary Figure 1.** Adding the relevant GWAS summary statistics to user profile.

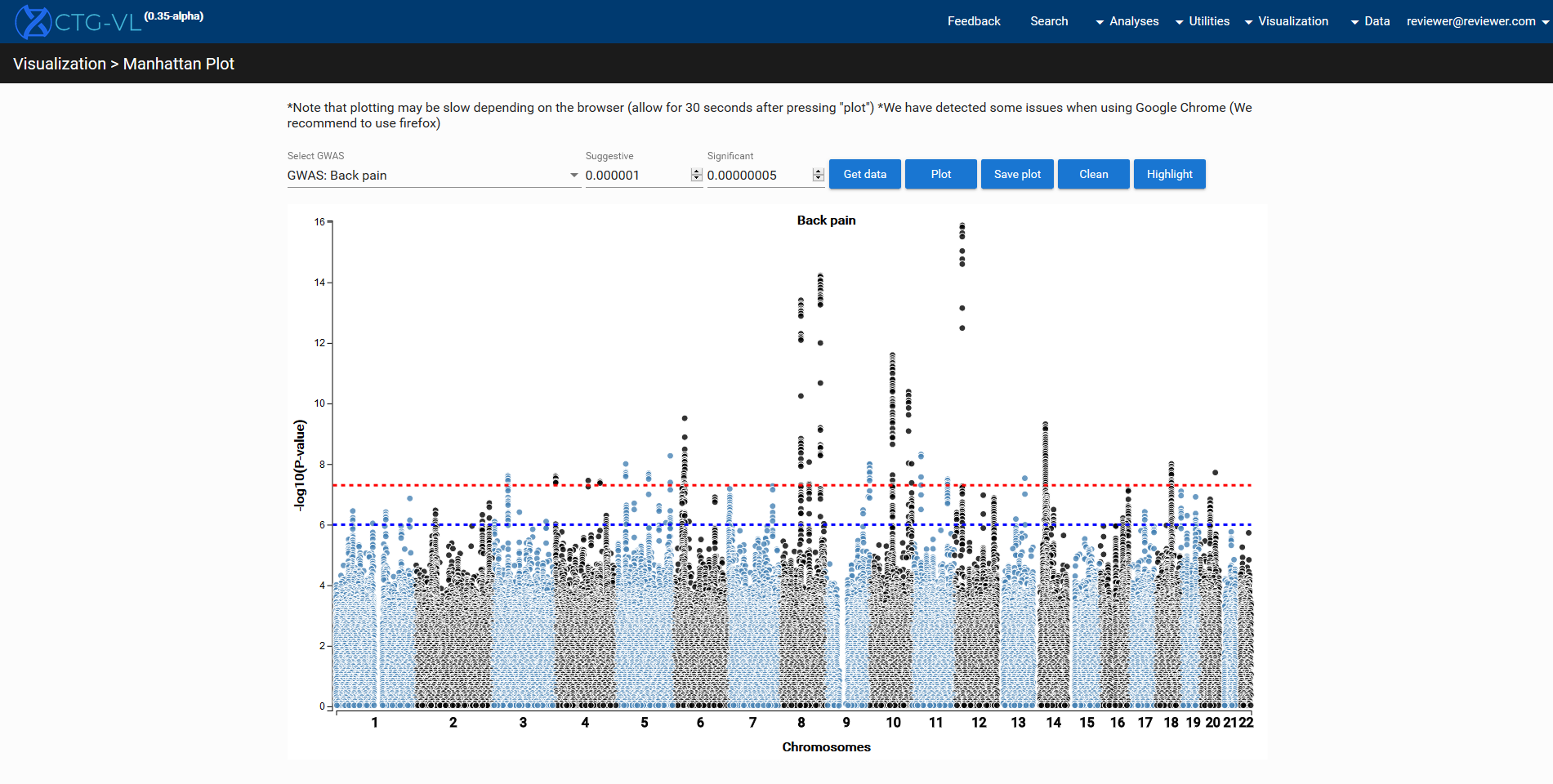

**Supplementary Figure 2.** Generating the Manhattan plot for back pain.

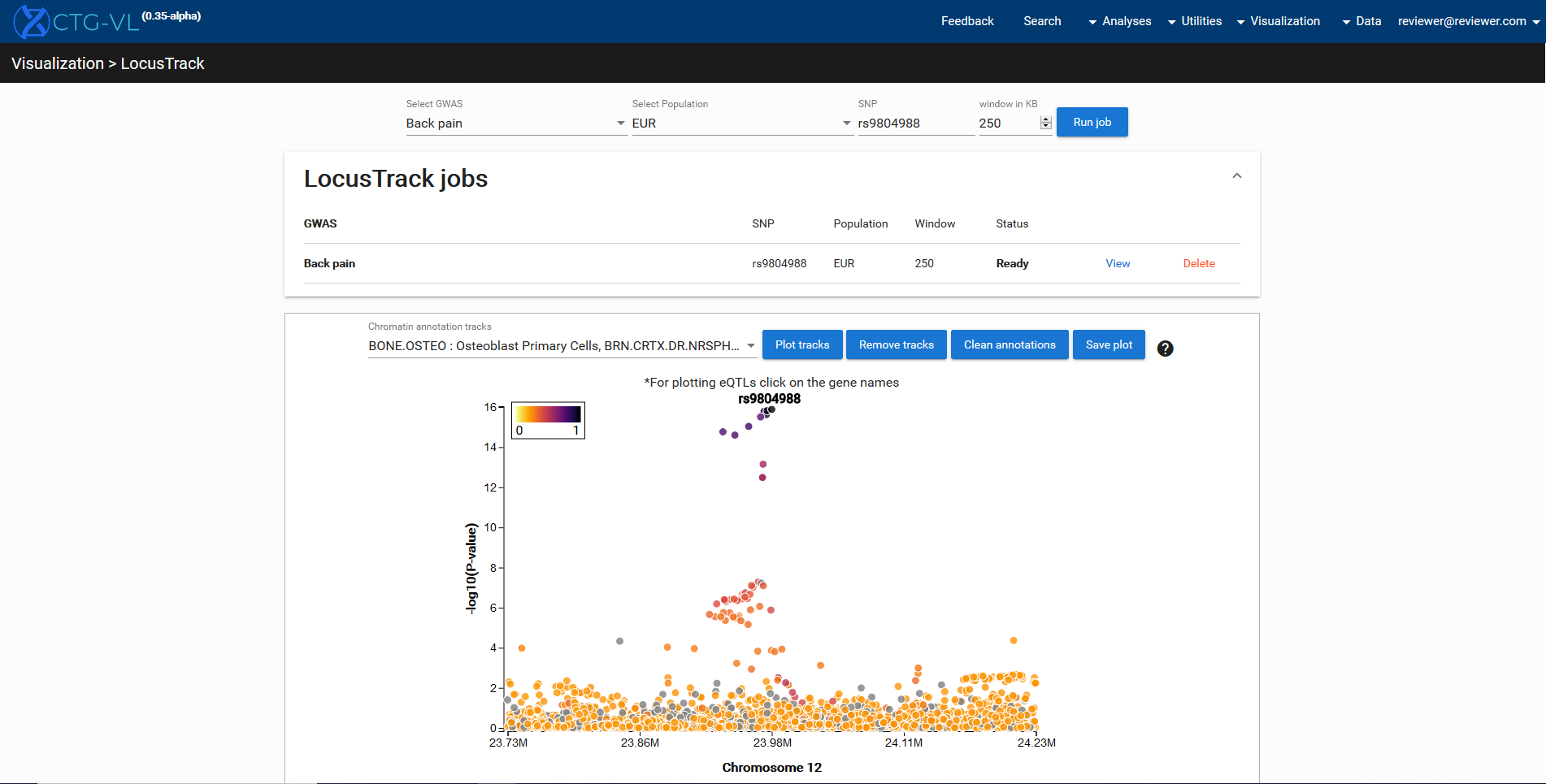

**Supplementary Figure 3.** Generating LocusTrack plot for the top GWAS hit.

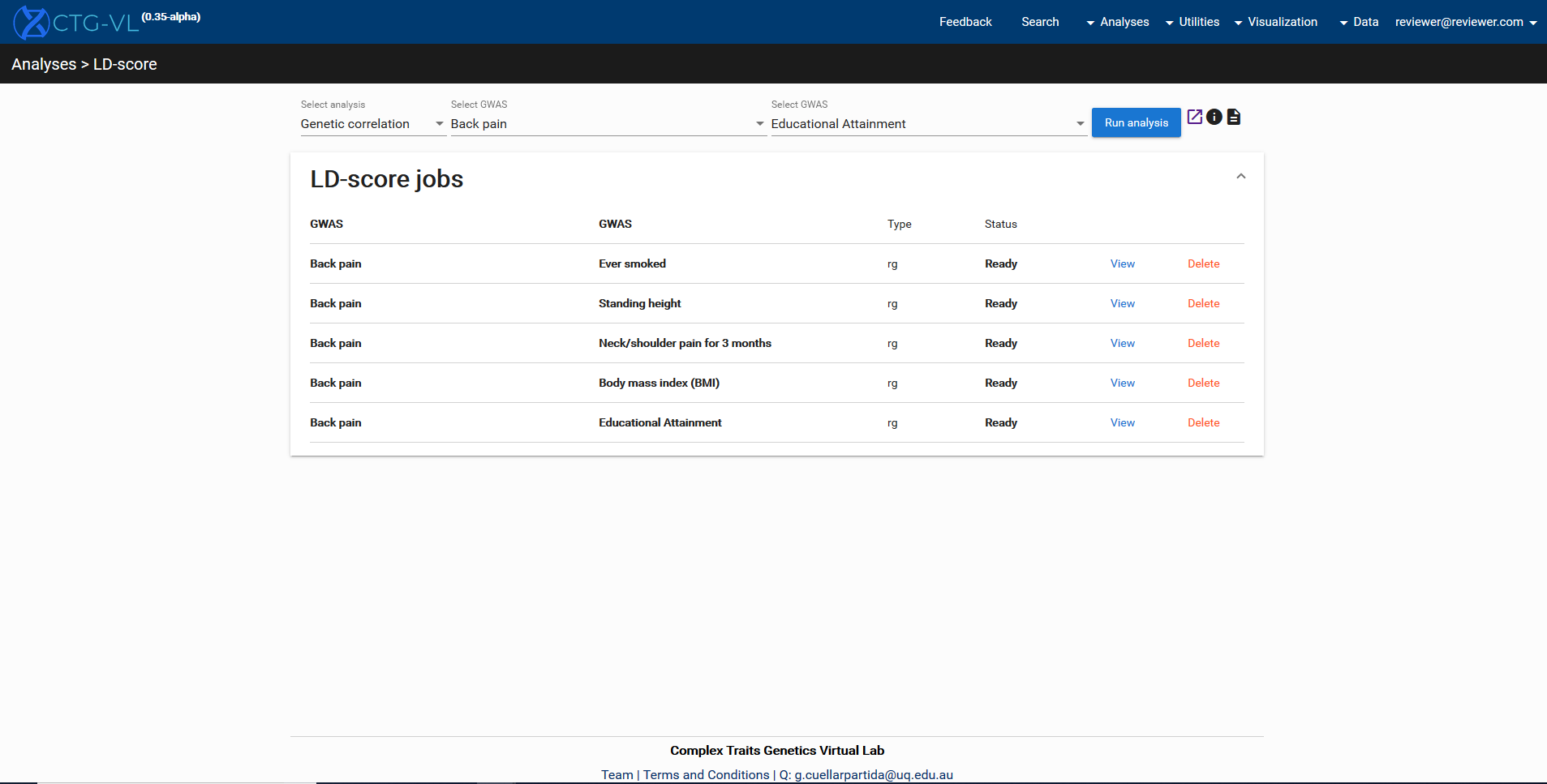

**Supplementary Figure 4.** Calculating genetic correlations with the relevant traits using LD-score regression.

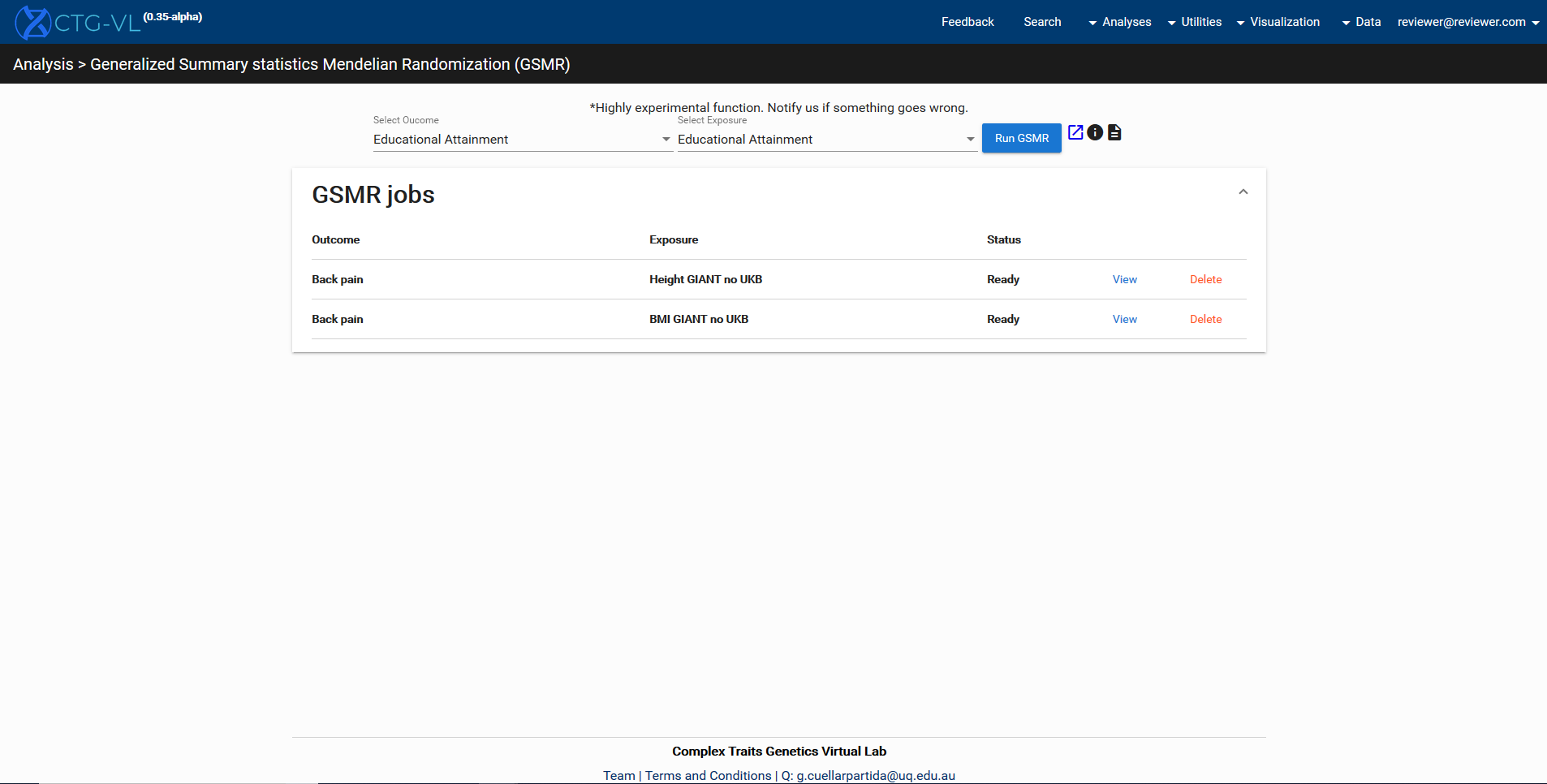

**Supplementary Figure 5.** Performing GSMR analyses to assess the causal relationship of BMI and height on back pain.

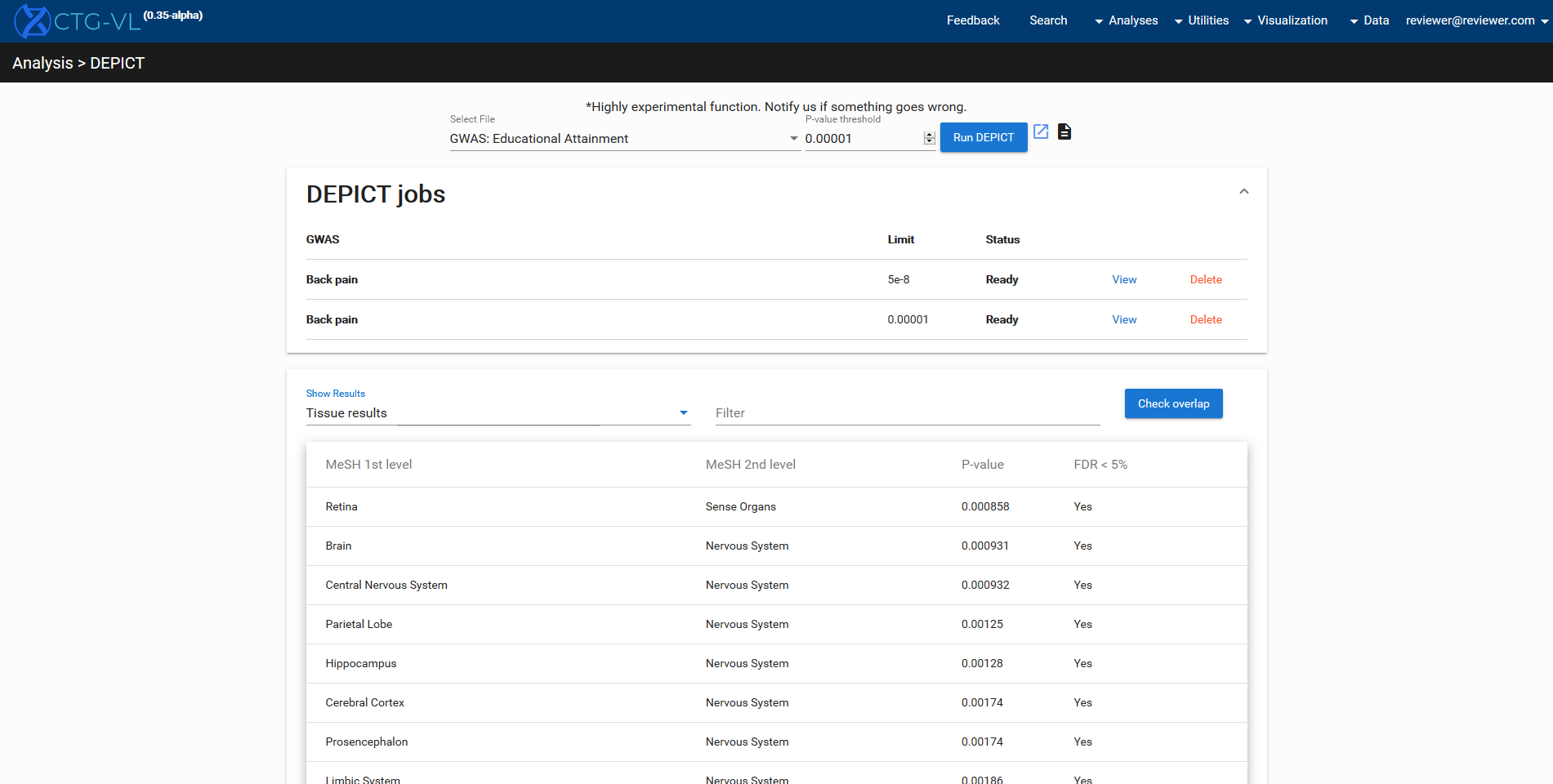

**Supplementary Figure 6.** Performing DEPICT analyses to identify tissues and biological pathways.

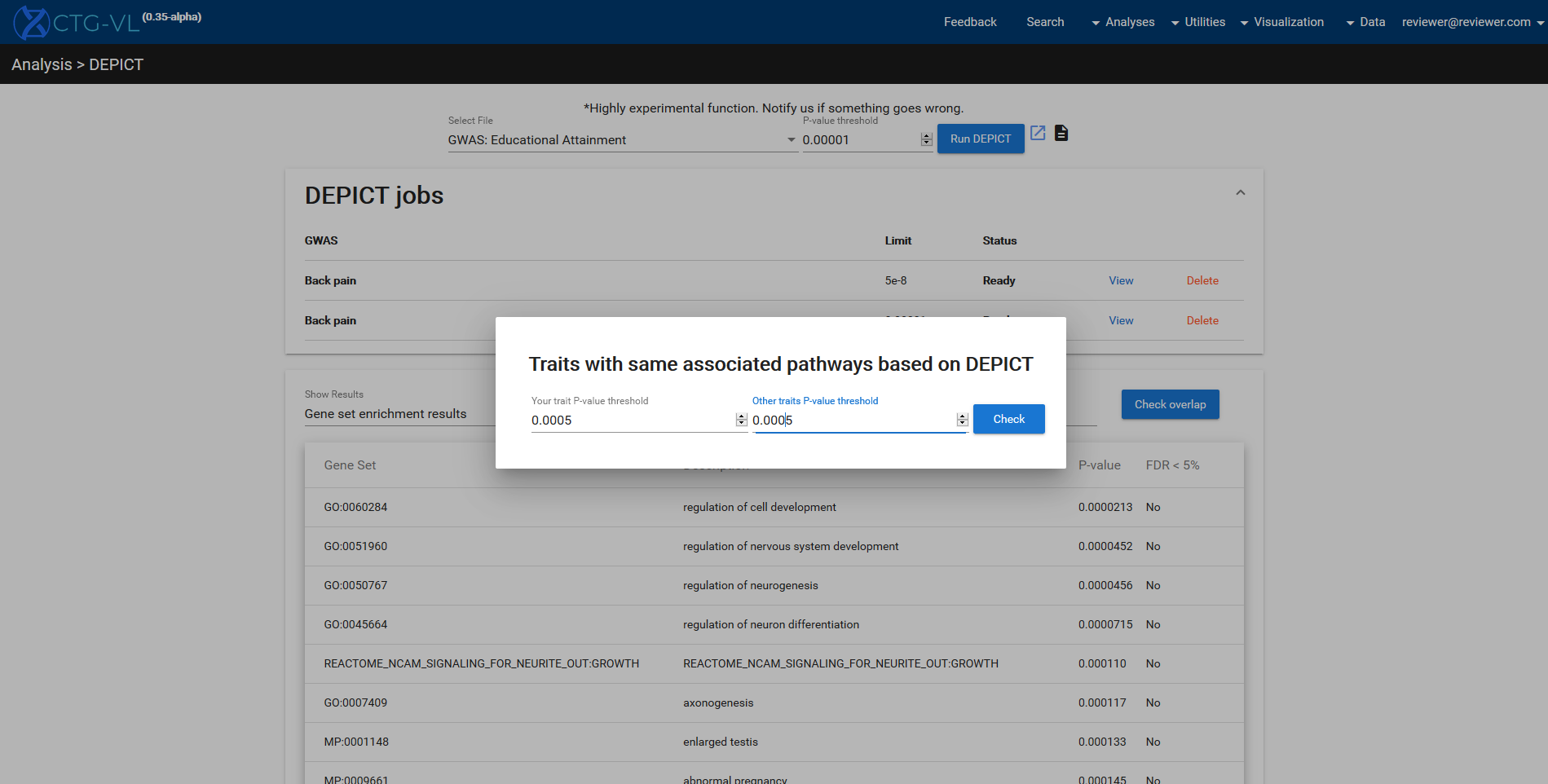

**Supplementary Figure 7.** Using the ‘check overlap’ function to identify other traits associated with the same biological pathways.

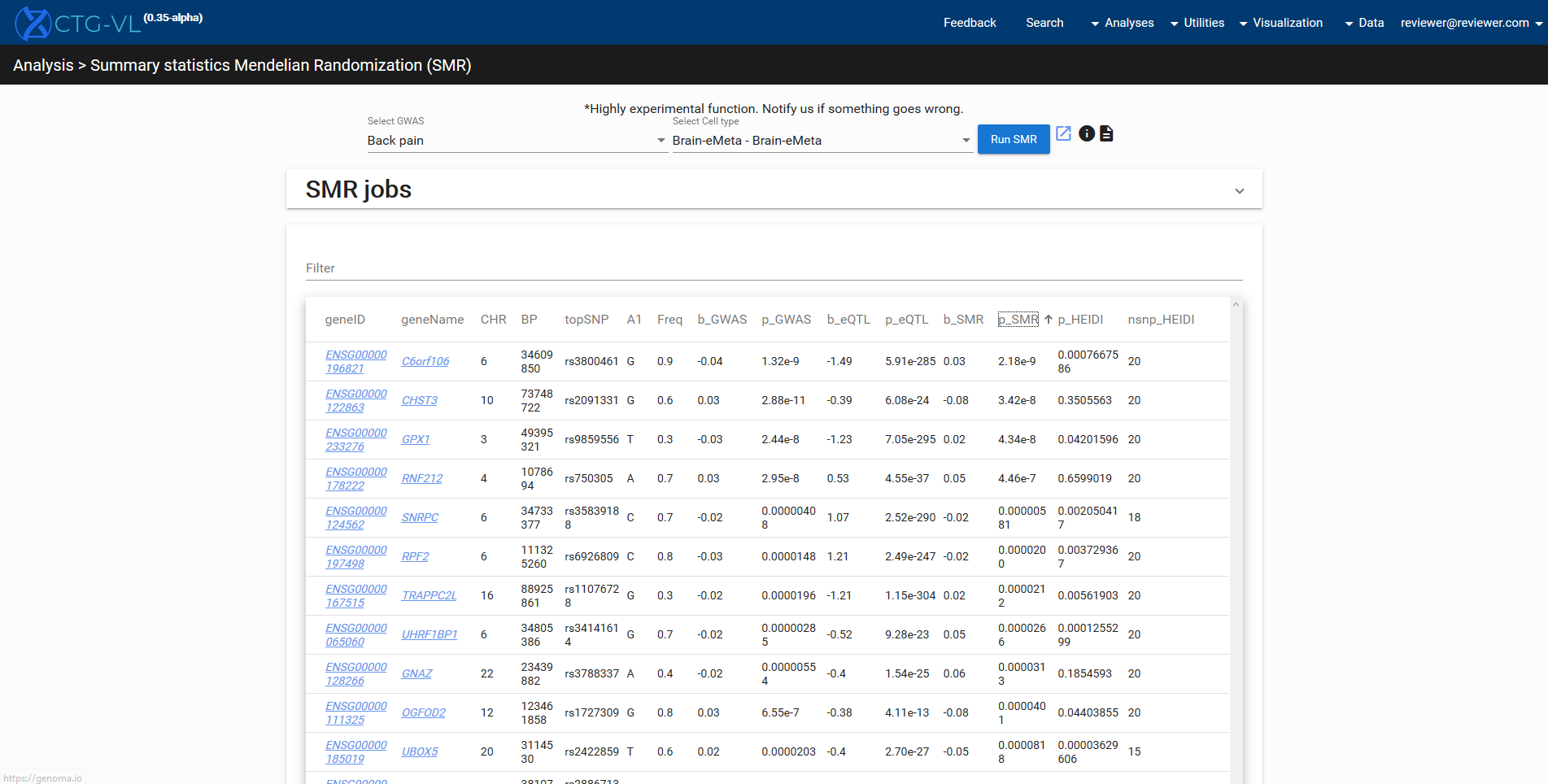

**Supplementary Figure 8.** Assessing causality between gene expression in brain tissues and back pain using SMR.

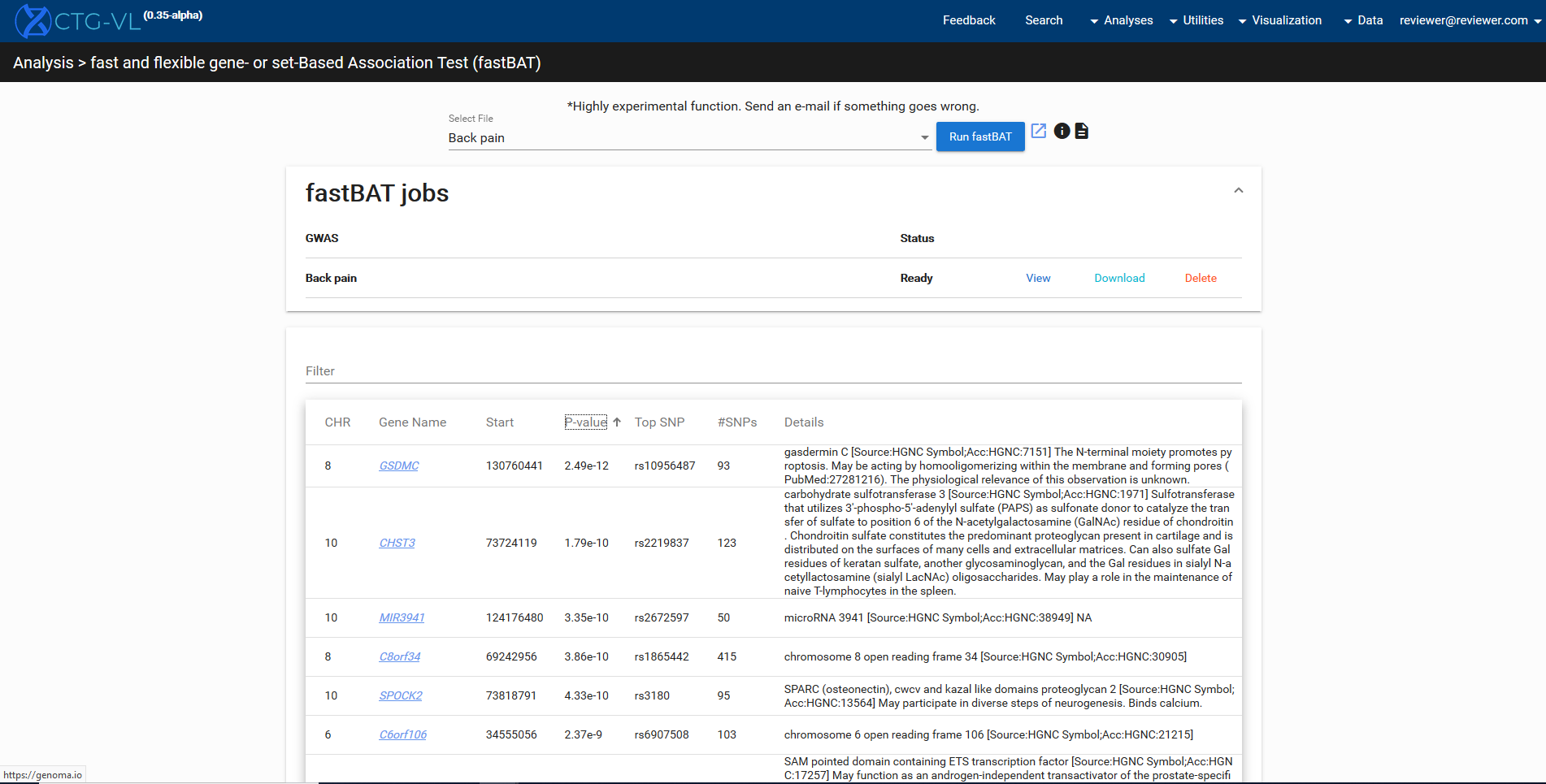

**Supplementary Figure 9.** Using fastBAT to perform gene-based association tests.
